# Sympathetic innervation regulates metabolic flexibility of skeletal muscle

**DOI:** 10.64898/2026.02.02.703364

**Authors:** Jordan Owyoung, HaoMin Sima, Junwon Heo, Tess Klugherz, Tina Tian, Brittney Ward, Nikki Boon, Garrett W. Cooper, Andrew L. Hong, Jarrod Call, Patricia J Ward

**Affiliations:** Department of Cell Biology, Emory University School of Medicine, Atlanta, GA; Department of Physiology and Pharmacology, College of Veterinary Medicine, University of Georgia, Athens, GA; Department of Pediatrics, Emory University School of Medicine, Atlanta, GA; Aflac Cancer and Blood Disorders Center, Children’s Healthcare of Atlanta, Atlanta, GA; Winship Cancer Institute, Emory University School of Medicine, Atlanta, GA; Regenerative Biosciences Center, University of Georgia, Athens, GA

**Keywords:** Sympathetic nervous system, skeletal muscle, B2-adrenergic receptor, mitochondria, CPT1

## Abstract

The sympathetic nervous system (SNS) is recognized for its role in the physiological regulation of organs, such as heart, vasculature and lungs, and has emerged as a potential player in skeletal muscle metabolic and neuromuscular junction (NMJ) health. However, the mechanism through which SNS signaling influences skeletal muscle function and adaptation to exercise remains unclear. Using molecular, electrophysiological, immunohistochemical, and high-resolution respirometry techniques, we tested the role of sympathetic innervation to skeletal muscle in response to exercise. Our findings reveal that sympathetic denervation disrupts the NMJ, reducing motor and sympathetic receptor expression, with concomitant deficits in skeletal muscle function. Mechanistically, these deficits are linked to diminished CPT1 enzyme activity, which impairs long-chain fatty acid-mediated oxidation in skeletal muscle mitochondria. These findings reveal a key role for sympathetic innervation in maintaining mitochondrial metabolic function and by extension, skeletal muscle performance, offering novel insight into the interplay between the SNS, exercise, and muscle mitochondria.

## Introduction

The role of sympathetic nervous system (SNS) is poorly understood in the context of skeletal muscle biology. The SNS is a branch of the autonomic nervous system that helps to maintain bodily homeostasis. SNS activity promotes pupil dilation, increased heart rate, and enhanced muscle performance—responses to stressful stimuli that give the SNS its nickname, “the fight or flight response”^[1, 2]^. The SNS has long been shown to innervate and control the function of organs, such as the heart, vasculature, and lungs, but much less is known about its innervation in skeletal muscle^[3-7]^. Recent studies have shown that sympathetic neurons likely send signals to muscle cells through receptors at the neuromuscular junction (NMJ)^[4, 8, 9]^. Dual innervation of NMJs by sympathetic and motor neurons was proposed in the early 1900s; however, it was not until 2016 that there was functional data to support SNS innervation in skeletal muscle^[8, 10-12]^.

Paravertebral sympathetic neurons innervate skeletal muscle and signal through release of norepinephrine (noradrenaline) neurotransmitters to adrenergic receptors in the NMJs^[4, 13, 14]^. In comparison, motor neurons also innervate skeletal muscle but use acetylcholine signaling to acetylcholine receptors (AchR) in the NMJ. This dual signaling sparks speculation on whether the SNS influences motor neuron signaling in the muscle. Khan *et al*. showed that a chemical SNS ablation results in decreased AchR concentration, indicating a possibility of crosstalk between sympathetic and motor neurons at the NMJ ^[8]^. While we know that the SNS is important for NMJ maintenance and function, how this correlates to skeletal muscle function or health is unknown.

Adrenergic receptors are a type of G-coupled receptor that bind catecholamines to elicit cellular signaling cascades^[15]^ and are the primary receptors of the SNS. There are two main types of adrenergic receptors (α and β) with various subtypes of each^[15, 16]^. In skeletal muscle, many sympathetic effects are mediated by β2-adgrenergic receptors (β2ARs)^[17]^. Sympathetic neurons signal to muscle cells through β2ARs, which are concentrated within the NMJ and selectively bind norepinephrine from the sympathetic terminals and epinephrine from the adrenal medulla. While physiological responses from adrenergic stimulation is tissue-specific, importantly in skeletal muscle, activation of β2ARs may increase muscle protein synthesis, inhibits muscle atrophy, and promotes mitochondrial biogenesis^[15, 18, 19]^. For example, β2AR agonism has been shown to attenuate muscle atrophy in mouse models of muscular dystrophy^[20-22]^. In the bodybuilding and fitness community, chronic use of β2AR agonists (sports doping) is common among athletes to induce skeletal muscle hypertrophy, supporting the notion that β2AR activation promotes muscle protein [23] synthesis^[15, 20]^. In mice, exercise has been shown to both increase β2AR concentrations in slow-twitch skeletal muscle and lower resting SNS activity^[24-27]^. Additionally, heart failure (HF) patients typically present with autonomic dysfunction caused by chronic overactivation of the SNS^[28, 29]^. Exercise has been shown to decrease muscle sympathetic nerve activity and norepinephrine output in heart failure patients^[28]^. While evidence supports that exercise transcriptionally regulates β2AR, it remains unknown whether exercise exerts beneficial skeletal muscle outcomes through primarily motor signaling or whether SNS signaling plays a major role^[27]^.

β2AR activation via exercise or the agonist formoterol increases levels of peroxisomal proliferator-activated γ coactivator 1α (PGC1a), a mitochondrial biogenesis transcriptional regulator^[30, 31]^. Thus, activation of β2AR by exercise potentially regulates mitochondrial biogenesis and function. In fact, data suggest changes in mitochondrial content, function, and substrate usage in response to exercise^[30, 32, 33]^.These changes can occur after as little as a single bout of endurance exercise^[32, 34]^. Mitochondria are particularly important to skeletal muscle health as they are critical for supplying ATP needed for muscle contraction, maintaining and regulating muscle metabolism, and NMJ function^[35-37]^. Frequently, mitochondrial diseases have detrimental effects on motor function due to the high density of mitochondria in skeletal muscle^[35, 37-39]^. Further, patients with spinal cord injury who are affected by skeletal muscle paralysis (but not muscle denervation) are at a much higher risk for Type II diabetes and obesity, two disorders that are associated with mitochondrial dysfunction, indicating the importance of mitochondrial health for not only skeletal muscle, but also for whole-body health^[40-43]^.

Research on exercise and muscle hypertrophy has shown the importance of exercise on muscle health^[44, 45]^. Endurance exercise, defined as any exercise that increases the capacity of the aerobic system, increases the size of the NMJ in a variety of hindlimb muscles, including the gastrocnemius and soleus muscles.^[46, 47]^ In aged mice, exercise has been shown to reverse and inhibit the denervation and loss of size of NMJs commonly seen in aged muscles^[46, 48-50]^. While exercise is established as beneficial for skeletal muscle, because the NMJ receives synaptic signaling from both motor and sympathetic neurons, the relative contribution of exercise through the sympathetic neuron at the NMJ remains to be resolved.

Our following data in mice support that SNS innervation is necessary for exercise-dependent changes in skeletal muscle and skeletal muscle mitochondrial function. To selectively ablate the sympathetic nervous system while preserving motor neuron innervation and function, we performed surgical lumbar sympathectomies on adult mice, resulting in regional (hindlimb) loss of sympathetic innervation. To evaluate the effect of exercise, subsets of mice were exposed to an exercise treadmill running regime. With this mouse model, we show that SNS innervation is necessary for exercise-induced functional improvements in skeletal muscle. We then show that exercise requires SNS innervation for mitochondrial substrate usage, which likely contributes to the functional changes we measured in the skeletal muscle. Thus, with our model, our findings support the functional importance of SNS innervation in skeletal muscle. These findings reveal a new fundamental biological function of the SNS and have implications for neural repair, axonal degeneration/regeneration, and aging fields as the SNS plays an important role in skeletal muscle mitochondrial function.

## Materials and Methods

### Animals

Adult wild-type C57B1/6 male and female mice (Jackson Laboratory Stock No. 000664) were used. Lights were automatically turned off at 7:00 and turned back on at 19:00 daily. All studies were approved by the Institutional Animal Care and Use Committee of Emory University and in compliance with National Institutes of Health Laboratory animal care and guidelines.

### Surgical sympathectomy

Mice were randomized into 1 of 4 groups: unexercised (unex), exercised (ex), sympathectomy (sympx) + unex, sympx + ex. For surgical sympx, animals were initially anesthetized with inhaled 3% isoflurane in 1L/min oxygen and then maintained at 2% isoflurane in 1L/min oxygen. The sympathetic ganglia were exposed by creating a small excision in the ventral abdominal wall and gently retracting the organs. Sympathetic ganglia were identified and the L2-L5 ganglia were visualized (control) or excised (sympathectomized)^[4, 51]^. The organs were returned to their original anatomical positions. The abdominal wall was closed with 5-0 absorbable suture, and the skin was closed with 5-0 non-absorbable suture. Mice were allowed to recover for 1 week before starting the aerobic exercise protocol.

### Exercise Protocols

Acute exercise: mice were subjected to a forced treadmill acute exercise protocol adapted from previously published mouse studies^[52-54]^. To maintain similar exposure and stress levels between the exercise and unexercised groups, unexercised mice were also brought to the treadmill room, but they were not run. Mice started with a 5-minute walking warm up at 10m/min. This was followed by 20 rounds of 30 seconds running + 60 seconds walking. The walk speed was held consistent at 10m/min. The first round run started at 12m/min, the second round at 15m/min, third at 18m/min, and fourth through twentieth rounds were at 21m/min. Bilateral gastrocnemius muscles were collected at 1 hour after the last running interval. See **Table 1** for exercise protocol.

Exercise training: mice were subjected to a forced treadmill exercise protocol. This protocol was adapted from previously published mouse studies^[26, 55, 56]^. Mice were exercised for 17 days: 5 days a week for 3 weeks and 2 days. Exercise occurred between 10:00 and 12:00 daily. Each training day started with a 5-minute warm-up at a velocity of 10 m/min. On the first day, mice ran for 20 minutes at a speed of 15m/min. Over the next 4 days, duration increased incrementally to 60 min while speed held constant at 15m/min. For the second week, mice ran for 60 min at a speed of 1m/min. For the third week, mice ran for 60 min at an increased speed of 17.5 m/min. The last two days mice ran for 60 min at an increased speed of 20m/min. See **Table 2** for exercise protocol.

### In-vivo gastrocnemius muscle evoked force, fatigue, and electromyography

We performed two in-vivo functional muscle assays in isoflurane anesthetized mice. Using a force transducer and custom Labview® software (National Instruments, 2024) for acquisition of electrophysiological data, we quantified electrically-evoked electromyographic activity in the form of compound muscle action potentials and force. First, a vertical skin incision was made in the hindlimb over the biceps femoris. The biceps femoris muscle was removed, and the gastrocnemius (GA) muscle and sciatic nerve were exposed. Stimulating electrodes were placed on each side of the sciatic nerve at approximately 1.5cm proximal to the GA muscle. To capture compound muscle action potentials (cMAPs), 2 fine wire recording electrodes were inserted into the GA muscle, 1 into the medial head and 1 into the lateral head. The subtalar joint was separated with scissors, preserving the attachment of the calconeal tendon to the calcaneus bone, and the calconeal tendon was then attached to the force transducer. Evoked muscle force and cMAPs were recorded in response to electrical stimulation of the sciatic nerve. The stimulus intensity began below motor threshold and was slowly increased until the cMAP and force output reached a plateau (maximum evoked force).

Fatigue assays were recorded in the same preparation. To achieve muscle fatigue, high frequency nerve stimulation was applied (100Hz and 60Hz) at 4 different stimulus intensities. Trial 1 stimulus intensity was set to 50% of the maximum evoked force. Trial 2 stimulus intensity was set to 80% of the maximum evoked force. Trial 3 stimulus intensity was set to 100% of the maximum evoked force. Trial 4 stimulus intensity was set to 120% of the maximum evoked force. These calculations were performed for each mouse. Each trial was 1000ms.

### RNA expression

Total RNA was extracted from GA muscle of acutely exercised and sedentary male and female mice using the RNeasy Fibrous Tissue Mini Kit (Qiagen) and cDNA was synthesized using SuperScript III First-Strand Synthesis Supermix kit (Thermo, USA). qPCR was performed using the QuantiTect SYBR Green PCR kit (Qiagen, USA). All samples were analyzed in ≥3 biological replicates. Primers were run in triplicates. Fold changes were calculated using the ΔΔCt method.

**Table 3.**
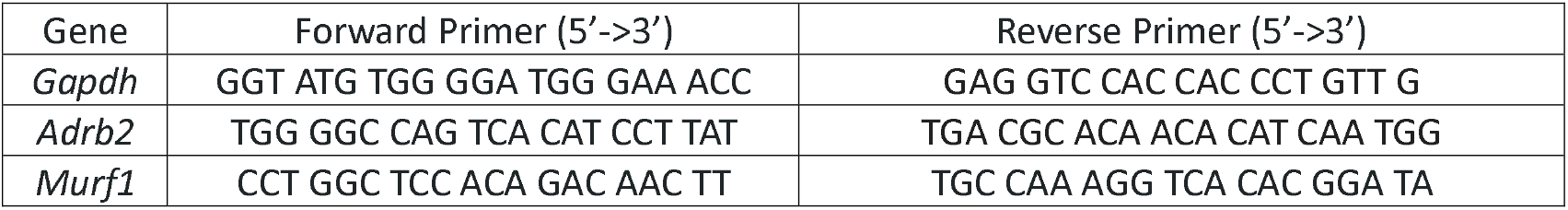
Primer sequences used for reverse transcription qualitative PCR.

### mRNA sequencing

Total RNA was extracted from mouse extensor digitorum longus (EDL) muscle using the RNeasy Fibrous Tissue Mini Kit (Qiagen) and quantified with the NanoDrop One spectrophotometer. For mechanistic studies, RNA was collected in biological duplicates from mice in the following treatment groups: unex, ex, sympx + unex, sympx + ex. Libraries were prepared using Illumina TruSeq. Samples were run with at least 20 million paired-end reads using Novoseq 6000 (Illumina) (Novogene, Sacramento, CA). Fastq files were aligned using HISAT2 to GRCm38.p3. Samples were then quantified using salmon through Illumina Dragen v3.7.5 on Amazon Web Services. Gene count files were converted to a counts matrix using tximport^[57]^. The counts matrix was used as input into DESeq2 to evaluate differential gene expression^[58]^. Normalized read counts matrices were used and input into principal component analysis to visualize clustering between samples using prcomp in base R[59]. Hierarchical clustering was performed using normalized gene expression counts. A distance matrix was computed based on 1 minus the Pearson correlation coefficient, and clustering was performed using the complete linkage method in base R. The resulting dendrogram was visualized using the ggdendrogram function from the ggplot2 and ggdendro packages in R. Significant genes with a LFC > |1| were used as input into gene ontology enrichment analysis using enrichGO and GO.db in the clusterProfiler package to identify genes in pathways that may be driving observed phenotypes. Versions used: R v4.2.2; R Studio 2022.07.1 Build 554; tximport v1.26.1, DESeq2 v1.38.3, ggplot v3.4.2, GO.db v3.16.1, ggdendro: v0.1.23, clusterProfiler 4.6.2.

### Immunofluorescent staining

Gastrocnemii muscles were dissected from isoflurane-anesthetized mice and post-fixed in 4% paraformaldehyde for 1 hour at room temperature. Muscles were then cryoprotected with 20% sucrose with 0.02% sodium azide overnight at 4°C. Muscles were placed in OCT, frozen, and longitudinally sectioned using a cryostat at -20°C. The 30 μm thick sections were placed onto charged slides.

Sections were permeabilized with 0.1% Triton®-X-100 (Fisher BioReagents, BP151-500) in 1X PBS then blocked for 2 hours at room temperature (RT) with 2% bovine serum albumin (BSA, Sigma-Aldrich, A2153), 0.1% Triton in 1X PBS. Slides were incubated with rabbit anti-β2AR (1:750, Abcam, Ab182136), or overnight at 4°C in a humidity chamber. After 3 x 10-minute washes with 1X PBS, secondary antibody goat anti-rabbit Alexa Fluor™ 488 (1:200, Invitrogen, A11008) was diluted in the blocking buffer and allowed to incubate at RT for 2 hours followed by 3 x 10-minute washes with 1X PBS. Following the washes, α-bungarotoxin (α-BTX) Alexa Fluor™ 555 (1:500, Invitrogen, B35451) was diluted in 1X PBS and placed on the slides for 30 minutes. Slides were washed 3 x 10-minutes with 1X PBS. After the slides were dried, they were mounted with Fluoro-Gel with Tris buffer and imaged on a Nikon Ti-E fluorescent microscope. A 60x oil objective was used with the following exposure times: TRITC 40 ms, Cy5 30 ms.

### Motor endplate area and quantification of AchR2 and β2AR fluorescent intensity at the NMJ

The motor endplate area (MEP) and relative expression of AchR2s on the post-synaptic terminal was determined by measuring the average fluorescence intensity of BTX and β2AR, respectively, at the NMJ. Images of NMJs were opened in FIJI, and a composite max projection was created. Each NMJ was outlined using the threshold feature, and the tracing was manually revised using the Wand Tool. The selected area was overlaid on either the AchR or β2AR signal to obtain mean intensity. Three additional areas were marked as designated background selections; these areas were obtained in regions surrounding the NMJ. Averages for the three background measurements were calculated. The ratio of fluorescence intensity of AchR or β2AR to background was calculated for each NMJ.

### Mitochondrial oxygen respiration

High-resolution respirometry was conducted on EDL muscle using an Oroboros Oxygraph-2K (Oroboros Instruments, Innsbruck, Austria) as previously described with a slight modification^[60]^. Experiments were carried out at 30°C in Buffer Z (105 mM K-MES, 30 mM KCl, 1 mM EGTA, 10 mM K2HPO4, 5 mM MgCl, 0.5 mg/mL bovine serum albumin; pH 7.1) supplemented with creatine (5 mM). Following the dissection of the EDL muscle from live mice under isofluorane, the tissues were chemically permeabilized with saponin (30 ug/ml), weighed (∼2mg), and placed into the chambers. The assay began with the addition of ADP (500 uM), followed by pyruvate/malate (PM; 1 mM/2 mM). Next, the inhibitor of pyruvate carrier, acyano-(1-phenylindol-3-yl)-acrylate, (UK5099; 1 uM) was added to block pyruvate entry, followed by palmitoyl-carnitine (Pal-Car; 40 uM) or octanoyl-carnitine (200 uM). Cytochrome c (10uM) was added to check the mitochondrial integrity, and then rotenone (ROT; 0.05 uM) was added to inhibit NADH-linked respiration. Finally, succinate was added to fuel complex II-linked respiration. All data were normalized to citrate synthase (CS) activity, an indirect marker of mitochondrial content.

### Assessment of mitochondrial enzyme activities

Mitochondrial enzyme activities (CS, β-hydroxyacyl-CoA dehydrogenase; β-HAD, and carnitine palmitoyltransferase I; CPT1) were measured spectrophotometrically using a spectrophotometer (Molecular Devices, San Jose, CA, USA). CS enzyme activity, an indirect mitochondrial content marker, was assessed as previously described^[61]^ by following the CoA-SH release from oxaloacetate and acetyl-CoA 5⍰, 5⍰-Dithiobis 2-nitrobenzoic acid (DTNB) to form TNB (OD: 412 nm). β-HAD activity was determined by incubating homogenate in a buffer containing 100 mM triethanolamine, 451 mM β-nicotinamide adenine dinucleotide, and 5 mM ethylenediaminetetraacetic acid (EDTA), as previously described^[62]^. Carnitine palmitoyltransferase I (CPT1) was assessed as previously described^[63]^ by capturing CoA-SH from palmitoyl-CoA via DTNB to form TNB (OD: 412 nm) in the Tris-HCl–DTNB buffer (116 mM Tris-HCl pH 8.0, 2.5 mM EDTA, 2 mM DTNB). All enzyme activities were normalized to CS enzyme activity.

### Statistical Analysis

Statistical analyses were carried out using Graphpad Prism. Data were analyzed either using students t-tests or two-way ANOVA followed with Tukey’s post-hoc analysis as indicated in each figure legend. Differences were considered significant if the calculated probability values were <0.05. For EMG and force parameters, principal component analysis (PCA) was performed using the Graphpad Prism PCA analysis function. Statistics were performed on the first and second principal components.

## Results

### Exercise transcriptionally regulates the muscle-specific sympathetic receptor

Previous studies have shown that exercise influences SNS neural activity^[24, 25]^. To evaluate if exercise directly influences the SNS molecularly, we acutely exercised wild type mice and determined RNA transcription levels of *Adrb2*, the gene that codes for β2AR, as well as peroxisome proliferator-activated receptor-γ coactivator 1α (*Pgc1α*), a gene downstream of the β2AR signaling pathway in muscle (Figure 1A). β2AR signaling pathways specifically target skeletal muscle hypertrophy pathways by upregulating muscle protein synthesis genes, downregulating muscle proteolysis genes, and increasing mitochondrial biogenesis (Figure 1B)^[15, 17, 18, 20, 30, 31]^. We found that *Adrb2* is transcriptionally upregulated 1 hour after acute exercise in both male and female mice (Figure 1C). Additionally, *Pgc1α* is transcriptionally upregulated 1 hour after acute exercise in both male and female mice (Figure 1D). We also evaluated transcription levels of muscle RING-finger protein-1 (*MuRF1/Trim63*); a gene involved in muscle remodeling and also downstream of β2AR signaling (Figure 1A). *MuRF1* is known to be upregulated directly after an acute bout of exercise and downregulated in response to long-term exercise^[64]^. We found that *MuRF1* is significantly upregulated 1 hour after acute exercise (Supplemental Figure 1). We also evaluated transcription levels in a skeletal muscle specific β2AR knock-out mouse model. In the β2AR knock-out, we found that there was no longer an increase in *MuRF1* after exercise, indicating that β2AR is necessary for exercised-induced molecular changes in skeletal muscle *MuRF1* (Supplemental Figure 1).

**Figure 1.**
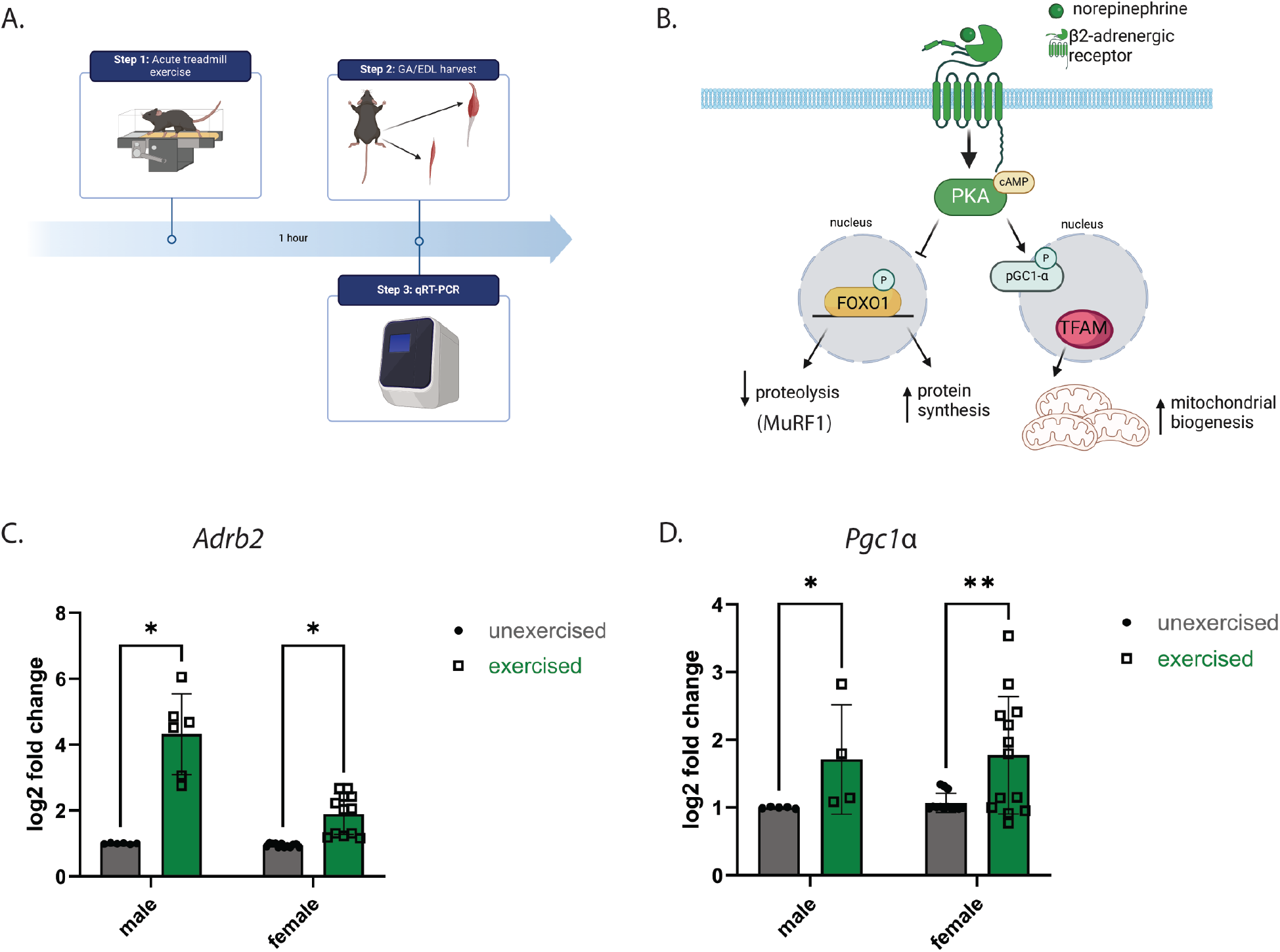
Exercise transcriptionally regulates the sympathetic receptor in skeletal muscle. **A.** Graphic showing timepoints for this set of experiments. GA muscle was collected and frozen 1 hour after acute exercise protocol. **B**. Graphical depiction of βeta-2 Adrenergic receptor (β2AR) downstream pathway. β2AR activation leads to activation of protein kinase A (PKA). This activation results in (1) inactivation of FOXO1 via hypermethylation, which drives an increase in protein synthesis genes and decrease in proteolysis genes and (2) activation of PGC1α, which results in an increase of mitochondrial biogenesis. **C-D**. Reverse transcription qualitative PCR results of Adrb2 (C) and Pgc1α (D) transcripts for male (left) and female (right) mice. Data shown as delta delta CT values normalized to an internal control (Gapdh). Treatments were compared using an unpaired t-test, *p < 0.05, **p < 0.01.

### Sympathetic innervation is required for NMJ morphology and exercise is sufficient to improve NMJ morphology

To evaluate NMJ morphology after sympathetic denervation, muscle sections were stained with BTX (Figure 2A-D). Following sympx, many NMJs lose their intricate, complex morphology compared to NMJs from naïve mice. We evaluated whether this change was significant by evaluating the total NMJ area. There was a trend for NMJs from sympathetically denervated muscle to be smaller compared to normally innervated mice (sympx + unex vs unex, p = 0.054). Exercise lead to increased NMJ size even in sympathetically denervated muscle (sympx + unex vs ex and sympx + unex vs sympx + ex p < 0.0001, p = 0.0191, Figure 2E). We also found that relative AchR fluorescent intensity is significantly reduced in sympathetically denervated muscle (sympx + unex vs unex NMJs (p < 0.001, Figure 2F).

**Figure 2.**
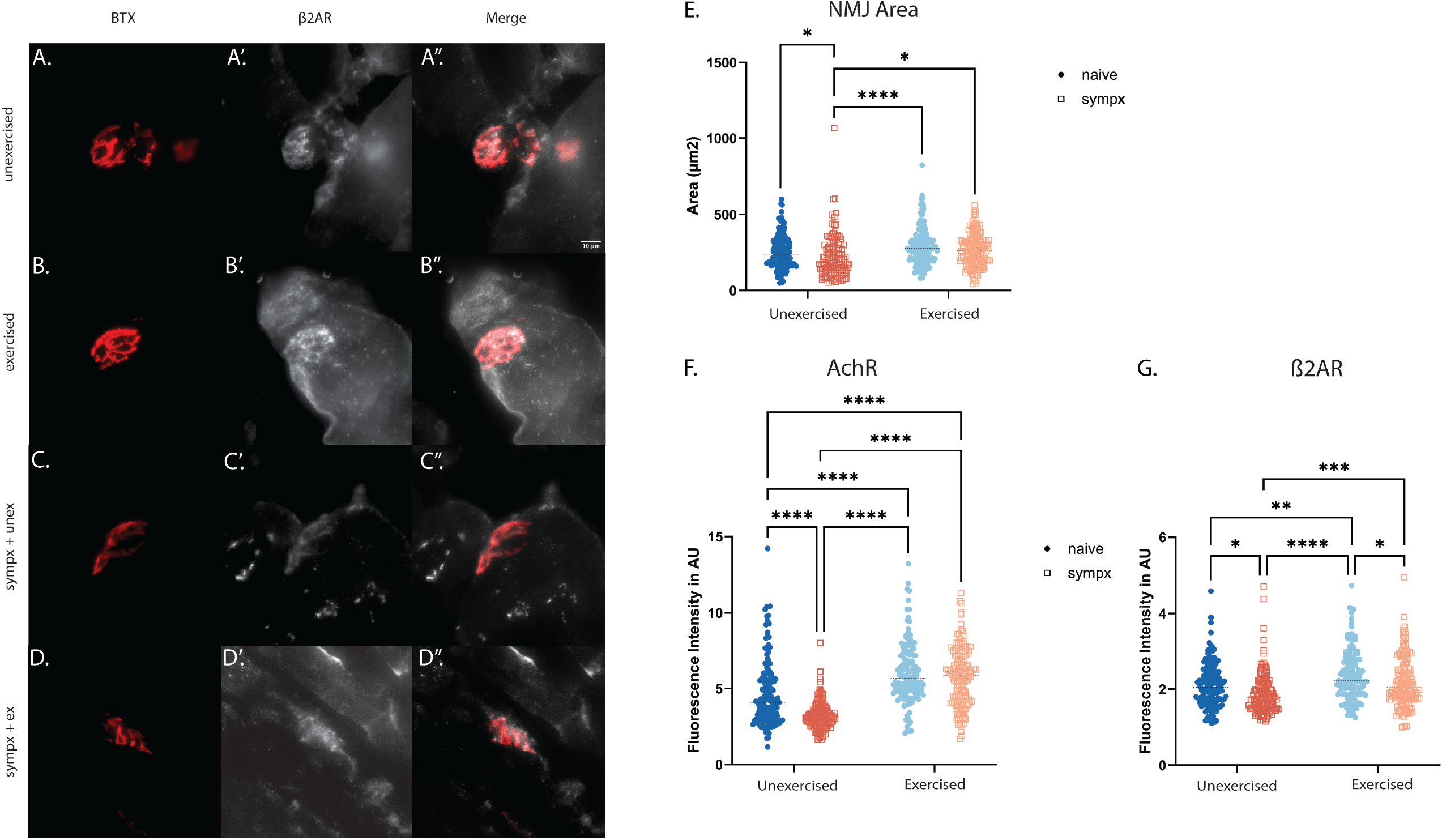
Sympathetic innervation regulates NMJ morphology and receptor density at the NMJ. A-D”. Images show representative NMJs from each treatment group(unex, ex, sympx + unex, sympx + ex) with bungarotoxin (BTX) (A-D) to visualize NMJs and anti-β2AR (β2AR) (A’-D’) and merged to visualize overlap (A”-D”). **E.** Quantification of NMJ area in GA muscle in unex, ex, sympx + unex, sympx + ex mice. **F**. Quantification of BTX (AchR) fluorescent intensity at the NMJ in GA muscle in unex, ex, sympx + unex, sympx + ex mice. Units shown in arbitrary units (AU). **G**. Quantification of β2AR fluorescent intensity at the NMJ in GA muscle in unex, ex, sympx + unex, sympx + ex mice. Units shown in arbitrary units (AU). A 2-way ANOVA with Tukey’s post-hoc analysis was performed for all quantifications, ****p < 0.0001, ***p < 0.001, **p < 0.01, *p < 0.05.

### Sympathetic denervation reduces neuromuscular junction β2AR fluorescent intensity and exercise is insufficient to recover it

Since AchR content at the NMJ was altered with sympathectomy, we were curious if the sympathetic receptor (β2AR) content followed the same pattern (Figure 2A’-D’). We found that there is a significant increase of β2AR fluorescent intensity at the NMJ in muscle in response to exercise (p = 0.001, Figure 2G). However, that increase is prevented with sympathectomy; both sympx + unex and sympx + ex have a significantly less β2AR fluorescent intensity compared to ex (p < 0.0001, p = 0.01) and are statistically similar to unex (Figure 2G). NMJs from sympathetically denervated muscle has significantly lower β2AR fluorescent intensity compared to all other groups. While sympx + ex is significantly increased compared to sympx + unex (p = 0.0007), it is still significantly decreased compared to ex (p = 0.01) (Figure 2F). Thus, exercise does not fully rescue the decrease in relative β2AR.

### Sympathetic innervation alters skeletal muscle function and exercise is insufficient to fully recover function

Because β2AR and AchR density levels were decreased in sympathectomized NMJs, we wanted to know whether this affected the function of the muscles even though motor neuron innervation remained intact. EMG, force, and fatigue assays were performed on the GA muscle 24 hours after the last day of exercise (Figure 3A). Representative fatigue curves (Figure 3B, B’), twitch force curves (Figure 3C, C’), and EMG waves (Figure 3D, D’) are shown. Data from these assays were linearly reduced and visualized using PCA. Interestingly, we found that evoked, *in-vivo* peak isometric force, cMAPs, and fatigue were all impacted by sympathectomy (Figure 3E). Unex and ex clustered together to the left of the y-axis and sympx + unex and sympx + ex groups clustered together to the right of the y-axis (Figure 3B). Tukey’s post-hoc analysis on PC1 values, primarily driven by peak isometric force, determined that ex was significantly different than sympx + unex (p < 0.0001) and sympx + unex (p = 0.001) groups and unex was significantly different than sympx + unex (Supp 2C). PC2 values were primarily driven by fatigue and did not show statistical significance (Supp 2D). This data indicates that sympathetic innervation contributes to skeletal muscle function, and functional differences in response to exercise is lost in animals lacking sympathetic innervation.

**Figure 3.**
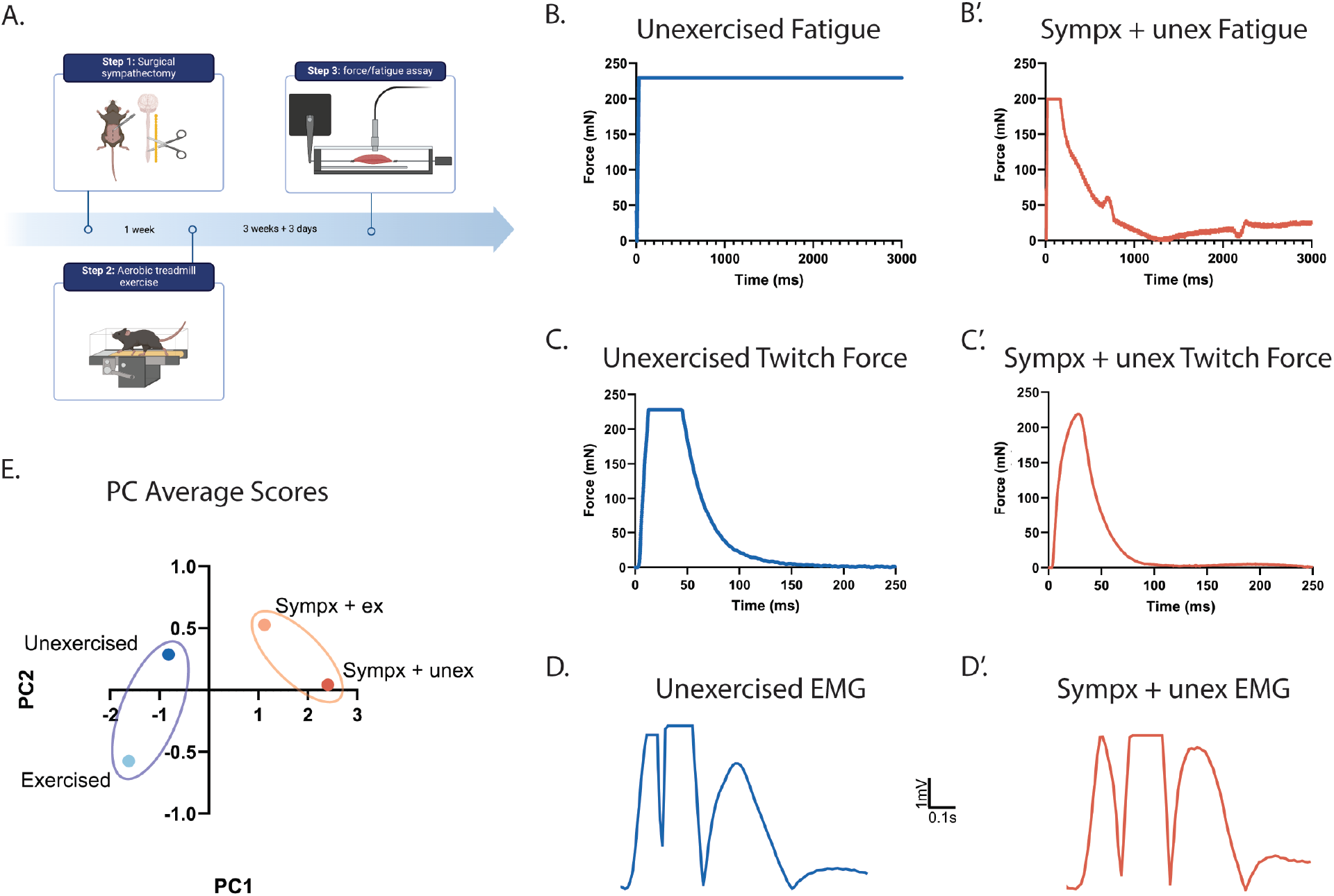
Sympathetic innervation alters skeletal muscle function and exercise is insufficient to recover function. **A.** Graphic illustration of experimental design. Mice were run using a treadmill exercise protocol 1 week after sympx surgery. Peak isometric force, fatigue, and EMG experiments were performed on the GA muscle 24 hours after the last day of exercise. **B**. Principal component analysis of peak isometric force, fatigue, and EMG data from GA muscle of unex, ex, sympx + unex, sympx + ex mice. Values averaged into one data point per treatment group. Ellipses are grouping statistically similar groups according to PCA1 values. **C-C’**. Example of fatigue curve in unexercised (C) and sympx + ex mice (C’). Recordings are shown as force over time in milliseconds. **D-D’**. Example of twitch force in unexercised (D) and sympx + ex (D’) mice. Recordings are shown as force over time in milliseconds. **E-E’**. Example of EMG curve in unexercised (E) and sympx + ex (E’) mice.

### Transcriptomic analysis of sympathetically denervated muscle

To better understand why peak isometric force is affected by sympathetic denervation, we evaluated the transcriptome of skeletal muscle. To do this, we performed bulk RNA sequencing on muscle from unex, ex, sympx + unex, and sympx + ex mice. Each treatment group was compared resulting in 6 pairwise comparisons (Supp Figure 3). Volcano plots show the differentially expressed genes (DEGs) between each comparison (Figure 4A-D). Upregulated DEGs for each comparison were grouped into signaling pathways using Gene Ontology (Figure 4A’-D’). Our sympx + unex vs unex data showed that upregulated genes are related to muscle movement and assembly suggesting that the SNS plays a role in skeletal muscle structure. Interestingly, the sympx + ex vs sympx + unex comparison has a large portion of upregulated genes that are part of the carboxylic acid transport and carbohydrate metabolism pathways. These pathways are not enriched in the DEGs from the ex vs unex comparison. This suggested that mitochondria could play a role in the functional changes in sympathetically denervated muscle (Figure 3).

**Figure 4.**
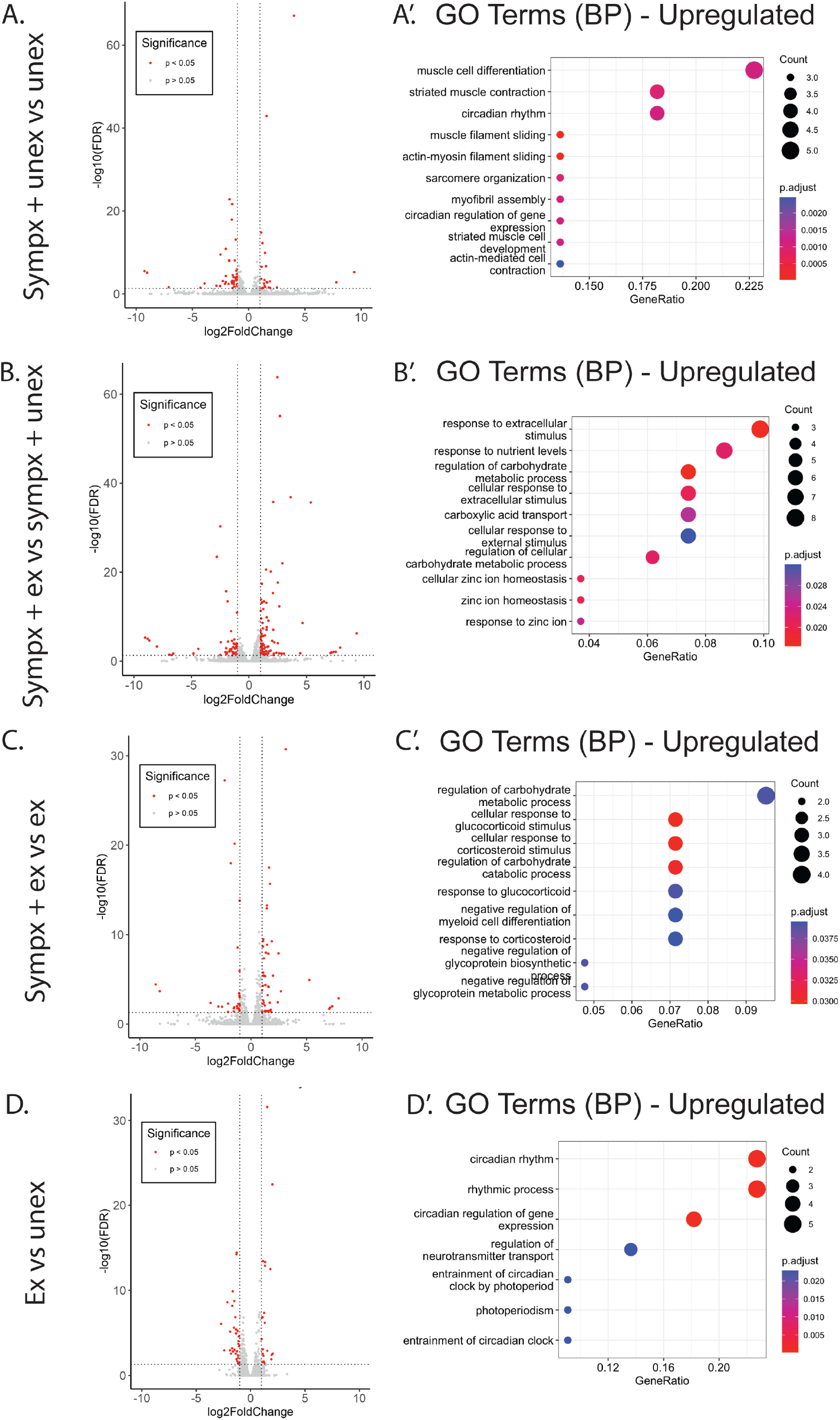
Transcriptomic analysis of sympathetically denervated muscle with and without exercise. **A-D.** RNA-sequencing results on EDL muscle visualized as volcano plots showing up (right) and down (left) regulated genes for each comparison: A. sympx + unex vs unex, B. sympx + ex vs sympx + unex, C. sympx + ex vs ex, D. Ex vs unex. **A’-D’**. Upregulated genes from each comparison grouped into Gene Ontology (GO) biological process (BP) pathways.

### Sympathetic denervation results in long-chain fatty acid B-oxidation deficits and exercise rescues this deficit

Because our RNA sequencing results showed an enrichment of DEGs related to mitochondrial metabolism, we were prompted to functionally test mitochondria in skeletal muscle. We performed all mitochondrial testing on muscle from the 4 treatment groups 24 hours after the last day of exercise (Figure 5A). To determine total mitochondrial content in the EDL muscle for each treatment group, we performed a CS activity assay. From this assay, we found no differences in mitochondrial content between any treatment group (Figure 5B). Because many of our DEGs are involved in mitochondrial metabolism, we wanted to determine whether there were differences in mitochondrial substrate preferences. We found that sympx + ex had significantly higher carbohydrate consumption (pyruvate and malate) compared to sympx + unex (p = 0.01) and ex groups (p = 0.05) (Figure 5C). Most interestingly from our mitochondrial substrate enzyme assay, we found that sympathectomy greatly decreases the ability of the mitochondria to oxidize long-chain fatty acids (i.e., palmitoylcarnitine). Both unex and ex groups had a much greater oxygen consumption rate from long chain fatty acids than sympx + unex groups (p = 0.04, p = 0.009) (Figure 5D). Exercise rescued this phenotype; sympx + ex and ex groups had statistically similar consumption rates while sympx + ex had a significantly increased consumption rate compared to sympx + unex group (p = 0.0024) (Figure 5D).

**Figure 5.**
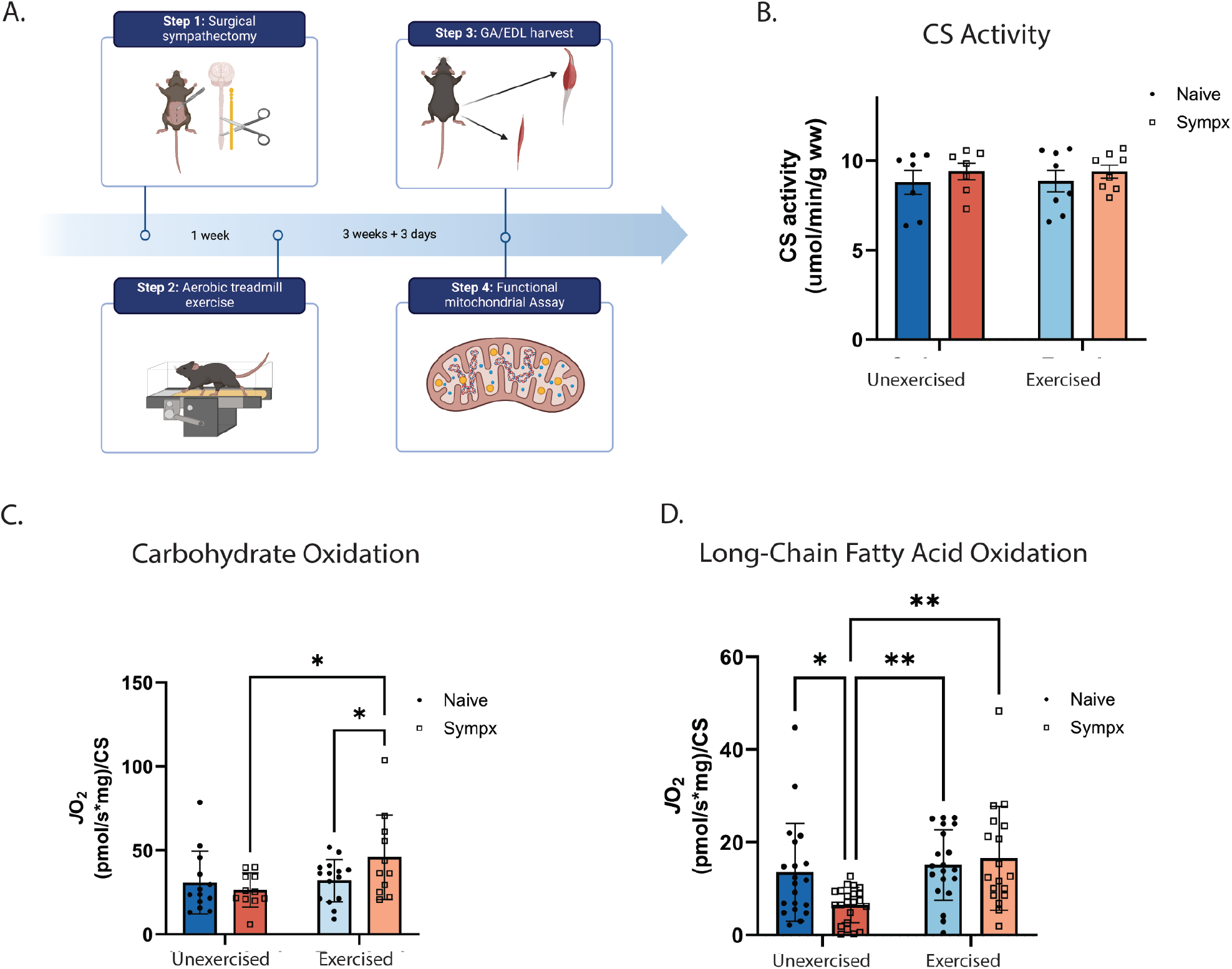
Sympathetic denervation results in long-chain fatty acid B-oxidation deficits. **A.** Graphic showing timepoints for this set of experiments. Mice were run using the aerobic exercise protocol 1 week after sympx surgery. Ex-vivo mitochondrial assays were performed on the EDL 24 hours after the last day of exercise. **B**. Citrate Synthesis enzyme activity representing levels of mitochondria in all four experimental groups. **C**. Muscles were fed pyruvate and malate to determine carbohydrate utilization differences. Sympx + ex showed a significantly increased consumption rate compared to ex and sympx + unex. **D**. Muscles were fed palmitoylcarnitine to determine long-chain fatty acid utilization differences. Sympx showed a significantly decreased consumption rate compared to unex, ex, and sympx + ex. A 2-way ANOVA with Tukey’s post-hoc analysis was performed for all quantifications, **p < 0.01, *p < 0.05.

### Sympathetic denervation contributes to CPT1 dysfunction which can be rescued with medium-chain fatty acid β-oxidation

We wanted to further probe components of mitochondrial respiration to determine where this deficit originated (Figure 6A). We started by evaluating complex I and II enzymatic functions, of which we found no changes between the four groups for complex I (Supp 4A)^[65]^. Sympx muscle also showed normal levels of complex II enzyme activity (Supp 4B). Next, we targeted an enzyme critical to the β-oxidation pathway itself, β-HAD, which we also found to be functionally similar in all four groups (Figure 6B). Lastly, we decided to evaluate the functionality of an enzyme critical for long-chain fatty acid oxidation specifically, carnitine palmitoyltransferase I (CPT1). CPT1 is a key enzyme in transporting long-chain fatty acids across the mitochondrial inner membrane for β-oxidation (Figure 6A). We found that the sympx + unex showed a significantly lower enzyme activity than ex (p = 0.009) and a trend towards significance compared to sympx + ex (p = 0.06) (Figure 6C), which agrees with our long-chain fatty acid oxygen consumption rate data (Figure 5). Seemingly, CPT1 enzyme activity correlates to long-chain fatty acid oxygen consumption rate. This supports CPT1 as a potential driver of mitochondrial substrate differences and therefore skeletal muscle deficits after sympx.

**Figure 6.**
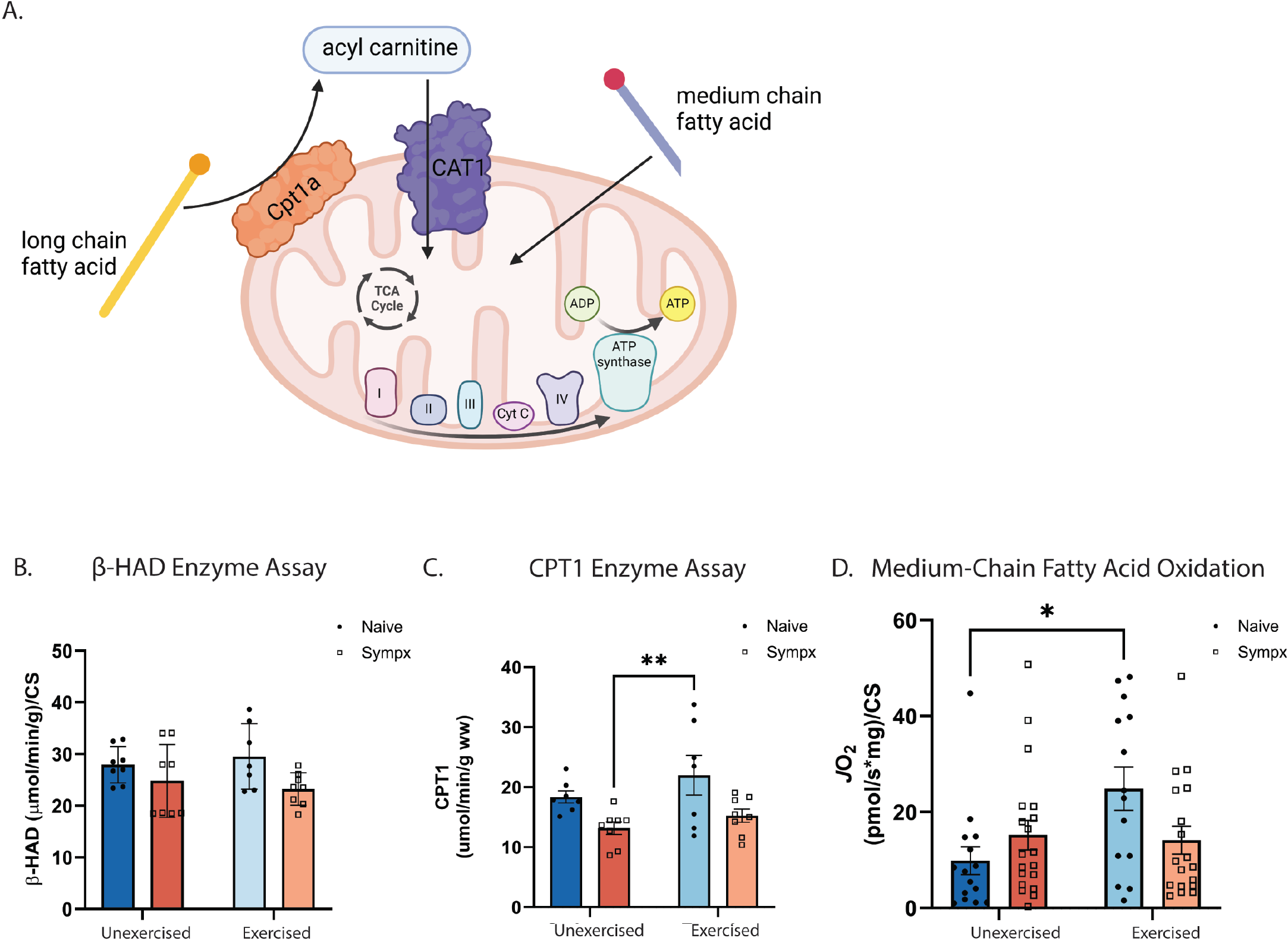
B-oxidation deficits from sympathetic denervation can be rescued with medium-chain fatty acid substrates. **A.** Graphical abstract highlighting the differences between long-chain fatty acid and medium-chain fatty acid transport into the mitochondria. Long-chain fatty acids require CPT1 and CAT1 to permeate the membrane while medium-chain fatty acids can permeate the mitochondrial membrane without enzymatic aid. **B**. 3-hydroxyacyl-CoA dehydratase (HACD1) assay showing similar levels of enzyme activity between the four groups. **C**. Carnitine palmitoyltransferase I (CPT1) assay showing decreased levels of enzyme activity in sympx + unex compared to ex groups. **D**. Muscles were fed octanoylcarnitine to determine medium-chain fatty acid utilization differences. Ex showed a significantly increased consumption rate compared to unex. A 2-way ANOVA with Tukey’s post-hoc analysis was performed for all quantifications, **p < 0.01, *p < 0.05.

Since sympathectomy resulted in a decrease in CPT1 function but no difference in β-HAD, we were prompted to test for oxygen consumption rates with medium-chain fatty acid substrates as CPT1 is not required for medium chain β-oxidation, but β-HAD is (Figure 6A). We tested using the medium-chain fatty acid octanoylcarnitine. Our data show that there is no longer a deficit in fatty acid consumption in the sympathetically denervated muscle compared to all other groups when fed medium chain fatty acid (Figure 6D). We next tested whole-body metabolic function of unex, ex, sympx + unex, and sympx + ex mice to probe whether these mitochondrial deficits were systemic. We found no overall difference in cage ambulation, respiratory exchange ratio, or carbohydrate and fat consumption (Supp 5A-D), indicating that the lumbar sympathectomy had local effects on the hindlimb muscles.

## Discussion

Although less understood than other targets, such as vasculature and cardiac cells, skeletal muscle is a direct target of SNS signaling. However, the role the SNS plays on skeletal muscle has remained ambiguous. Sympathetic innervation contributes to NMJ morphology, but how muscles physiologically respond to sympathetic denervation and subsequent NMJ dysregulation is unclear. Exercise has been shown to both increase sympathetic activity during and after an acute bout of training and decrease sympathetic activity after long-term training^[24, 25, 66]^. In addition, it has long been accepted that exercise results in muscle hypertrophy and increased muscle metabolism and health. Here, our data suggest, in part, a molecular connection via mitochondrial function between exercise, the SNS, and skeletal muscle health.

Previous data has supported the importance of exercise on the NMJ and skeletal muscle health; this has been shown by evaluating NMJ size and AchR density after exercise^[8, 46, 67]^. We also observed an increase in AchR density and NMJ area in the muscle after aerobic exercise. From previous data, we know that removing sympathetic innervation results in a decrease in AchR density and loss of morphology[8]. Our data also showed that sympathectomy resulted in a decrease of AChR, visual loss of complex morphology, and decreased motor endplate area. Endurance exercise is expected to increase NMJ area in mice, which we also observed in our sympathetically denervated muscle^[46]^. This rescue of synaptic area is likely due to the presence of functional motor innervation. Although systemic sympathetic activation is possible, e.g., norepinephrine/epinephrine release from adrenal glands, sympathetic denervation also resulted in less β2AR within the muscle synapse, and exercise was unable to rescue its expression. Furthermore, our evidence suggests that exercise induces transcriptional regulation of *Pgc1a, Murf1*, and *Adrb2*. As we no longer saw an increase in *Murf1* after exercise in the conditional knockout, B2AR-dependent changes in *Murf1* may underlie part of the muscle remodeling process. Altogether, this evidence supports the importance of direct SNS innervation in skeletal muscle and at the NMJ. ^[46]^

Of importance, sympathetic signaling plays a role in the control of muscle blood flow and maintenance of blood pressure^[66]^. It is possible our results are from a decrease in vascular tone affecting blood flow and thus a decrease in mitochondrial and muscle function. However, this is unlikely because exercise results in an increase in blood flow to the exercising limbs, and in a dog model of surgical sympathectomy, blood flow was actually increased in the sympathectomized limb^[68]^. Additionally, sympathetic neurons in the cervical and thoracic regions are preserved in our sympathectomy model, and these are the neurons that project to the heart and other vasculature^[1, 7]^. Thus, we believe our results are likely not due to decreased blood flow but rather from a lack of sympathetic input at the skeletal muscle itself. However, it is important to note that we did not monitor blood flow in these experiments, and future experiments to separate vascular effects from adrenergic signaling should be performed.

We found that skeletal muscle functional changes were accompanied by transcriptional changes as shown by our RNA-sequencing data. Following exercise, we found an increase in muscle myosin (*Myh7, Myh7B*), as expected with muscle hypertrophy. Additionally, we saw that many upregulated DEGs due to sympathetic denervation were involved in actin and myosin movement and assembly, indicating likely remodeling of the muscle. With exercise, sympathetically denervated muscle showed an increase in carbohydrate-related GO processes, which is in alignment with previous data showing increased carbohydrate substrates in response to an increased energy demand from training^[69]^.

Exercise promotes alterations to mitochondrial substate usage^[30, 32-34]^, thus, using high-resolution respirometry, we tested the oxygen consumption rates with various substrates in real-time (Figure 5). Exercise of sympathetically denervated muscle resulted in an even greater carbohydrate metabolism compared to naive exercised muscle, which is supported by our RNA-sequencing data. Of note, sympathetic denervation resulted in a significant decrease in oxygen consumption rates compared to all other groups when given palmitoyl-carnitine, indicating that long-chain fatty acid-mediated β-oxidation is decreased or impaired in sympathectomized muscles. Interestingly, exercise is able to rescue this metabolic defect.

Long-chain fatty acid β-oxidation begins in the mitochondrial matrix, where fatty acids are converted into acylcarnitine before transport into the inner mitochondrial membrane. There, acylcarnitine is oxidized to acyl-CoA, which is further oxidized in the mitochondrial matrix to produce molecular products such as NADH to be used in the electron transport chain. To test why there is a deficit in long-chain fatty acid β-oxidation, we first tested the enzyme activity of β-HAD, which is the third of four enzymes involved in converting acylcarnitine into acyl-CoA and has been shown to be a key factor in muscle cell growth^[70]^. Since there were no differences in β-HAD enzyme activity between the 4 groups, we next tested CPT1 activity, an enzyme involved in long-chain fatty acid transport across the inner mitochondrial membrane. This step is only necessary for long-chain fatty acids as short- and medium-chain fatty acids can permeate the membrane^[71]^. Sympathetically denervated muscle had decreased CPT1 activity, which could explain the decrease in long-chain fatty acid oxygen consumption rate. There were no changes in the mRNA levels of *Cpt1*, thus enzymatic activity changes in CPT1 are likely from differences in molecules that closely regulate it, such as malonyl-CoA or FATP1^[72, 73]^. Interestingly, medium-chain fatty acid-derived oxygen consumption rate, which does not rely on CPT1 activity, was not impaired by sympathetic denervation, and and also supports that long-chain fatty acid β- oxidation is being hindered at the CPT1 enzymatic step. Thus, our data shows that mitochondrial substrate differences after sympathectomy, which contributes to muscle dysfunction, are driven by a decrease in fatty acid oxidation caused by a functional decrease in CPT1. It is important to note that we used a treadmill-based running exercise protocol. Further studies should include a range of exercise types and lengths to assess their influence on skeletal muscles, mitochondria, and the SNS^[47, 74]^.

CPT1b (the main isoform of CPT1 found in skeletal muscle and heart tissue) deficiency has been shown to increase lipid content and whole body adiposity, which is often associated with Type II diabetes and obesity^[42]^. Further, skeletal muscle mitochondrial dysfunction is also related to the development of Type II diabetes and obesity^[40-43]^. It is highly likely that the development of these disorders is linked to CPT1 dysfunction, and an increase in CPT1 enzymatic activity enhances fatty acid oxidation to improve high-fat diet induced insulin resistance. Our data supports that changes in skeletal muscle mitochondrial respiration from sympathetic denervation is driven by a decrease in CPT1 activity. Therefore, promoting diets heavy in carbohydrates, short-, and medium-chain fatty acids, which do not require CPT1, could not only be beneficial for patients with sympathetic perturbations, but also for patients susceptible to developing Type II diabetes and obesity.

## Supplemental Materials and Methods

### Animals

Prior to muscle collection, mice were weighed on a milligram scale to determine body weight. Mouse tibias were collected after muscle collection and measured in millimeters (mm) using calipers.

### Adrb2 Knock-out mice

*Adrb2*^fl/fl^ mice were generously provided by Dr. Daniel Mucida at Rockefeller University^[75, 76]^. *Adrb2*^fl/fl^ mice were crossed to inducible HSA-cre mice (Jackson Laboratories Stock No: 025750). Mice were kept in reverse dark/light cycle rooms and allowed to habituate for 14 days before use. Lights were automatically turned off at 7:00 and turned back on at 19:00 daily. *Adrb2*^*f*l/fl^ with no cre recombinase (naïve) were used as controls for these experiments. Conditional knockout and control mice were treated with a 10mg/ml tamoxifen solution in sesame oil and dosed at 3.75µg/g of body weight for 6 days over 2 weeks (3 consecutive days in week 1 followed by 4 day recovery and additional 3 consecutive days in week 2). Mice were allowed to recover for 2 weeks prior to undergoing the acute exercise protocol. All studies were approved by the Institutional Animal Care and Use Committee of Emory University and in compliance with National Institutes of Health Laboratory animal care and guidelines.

### Indirect calorimetry (whole-body metabolism)

Promethion indirect calorimetry cages (Sable Systems International, Las Vegas, NV, USA) were used to determine energy metabolism in unex, ex, sympx + unex, and sympx + ex C57B1/6 mice. Mice were housed individually in metabolic cages with *ad libitum* access to water and food. They were acclimated to the cages for 20 hours before data collection began. Data collection continued for 72 hours in 5 minute increments. Energy expenditure (EE) per hour was calculated using a modified Weir equation [EE (kcal/hr) = 60 ⍰ (0.003941 × VO2+0.001106 × VCO2)], and the respiratory exchange ratio (RER) was calculated by the volume ratio of CO2 and O2. Cage activity was quantified in meters using the total distances calculated from Pythagoras’ theorem based on XY position coordinates recorded second by second.

## Supplemental Figures

**Supplemental Figure 1.**
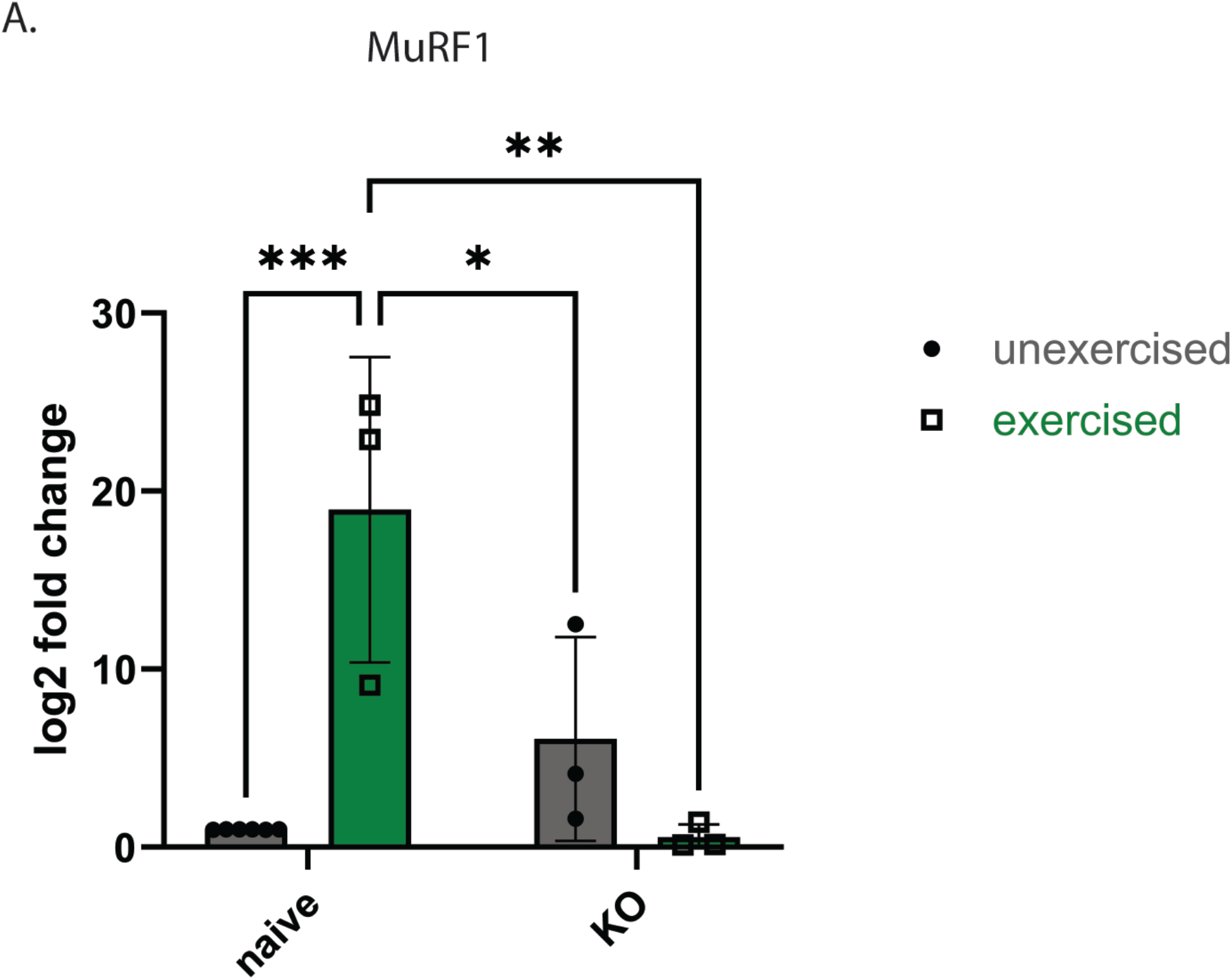
Exercise transcriptionally regulates MuRF1 in skeletal muscle. **A.** Reverse transcription quantitive PCR results of MuRF1 transcripts for naive (left) and β2AR knock-out (right) mice. For each comparison, mice were either unexercised (circle) or acutely exercised (square). Data shown as delta delta CT values normalized to an internal control (Gapdh). Treatments were compared using a 2-way ANOVA with Tukey’s post-hoc analysis, *p < 0.05, **p < 0.01, ***p < 0.001.

**Supplemental Figure 2.**
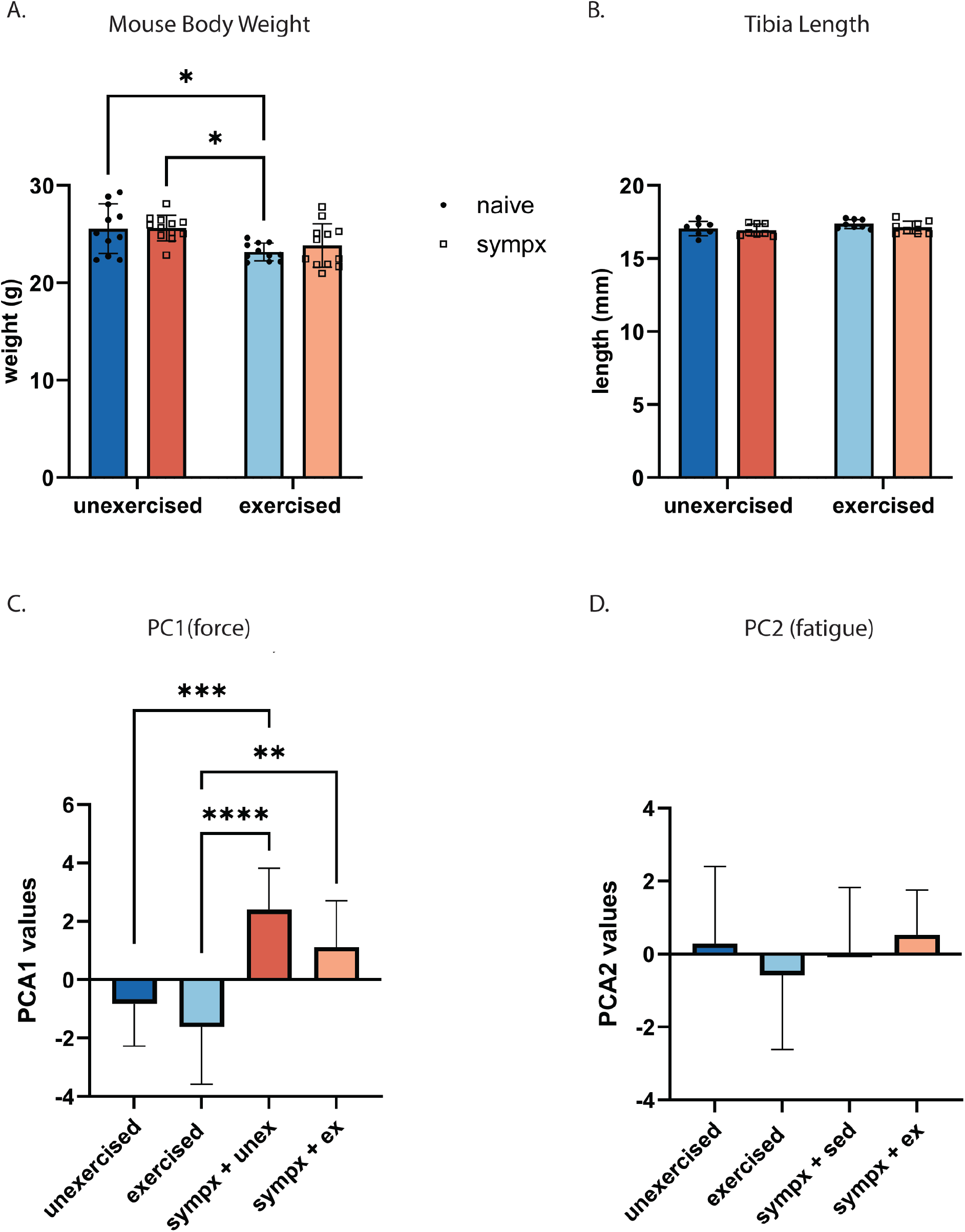
Skeletal muscle function is affected by sympathetic denervation. **A.** Body weights of experimental mice prior to force, fatigue, and EMG. **B**. Tibia lengths of experimental mice. **C-D**. PCA1 (C) and PCA2 (D) values of mice from all 4 experimental groups: unex, ex, sympx + unex, sympx + ex from PCA analysis (Figure 3E). A 2-way ANOVA with Tukey’s post-hoc analysis was performed for all quantifications, ****p < 0.0001, ***p < 0.001, **p < 0.01, *p < 0.05.

**Supplemental Figure 3.**
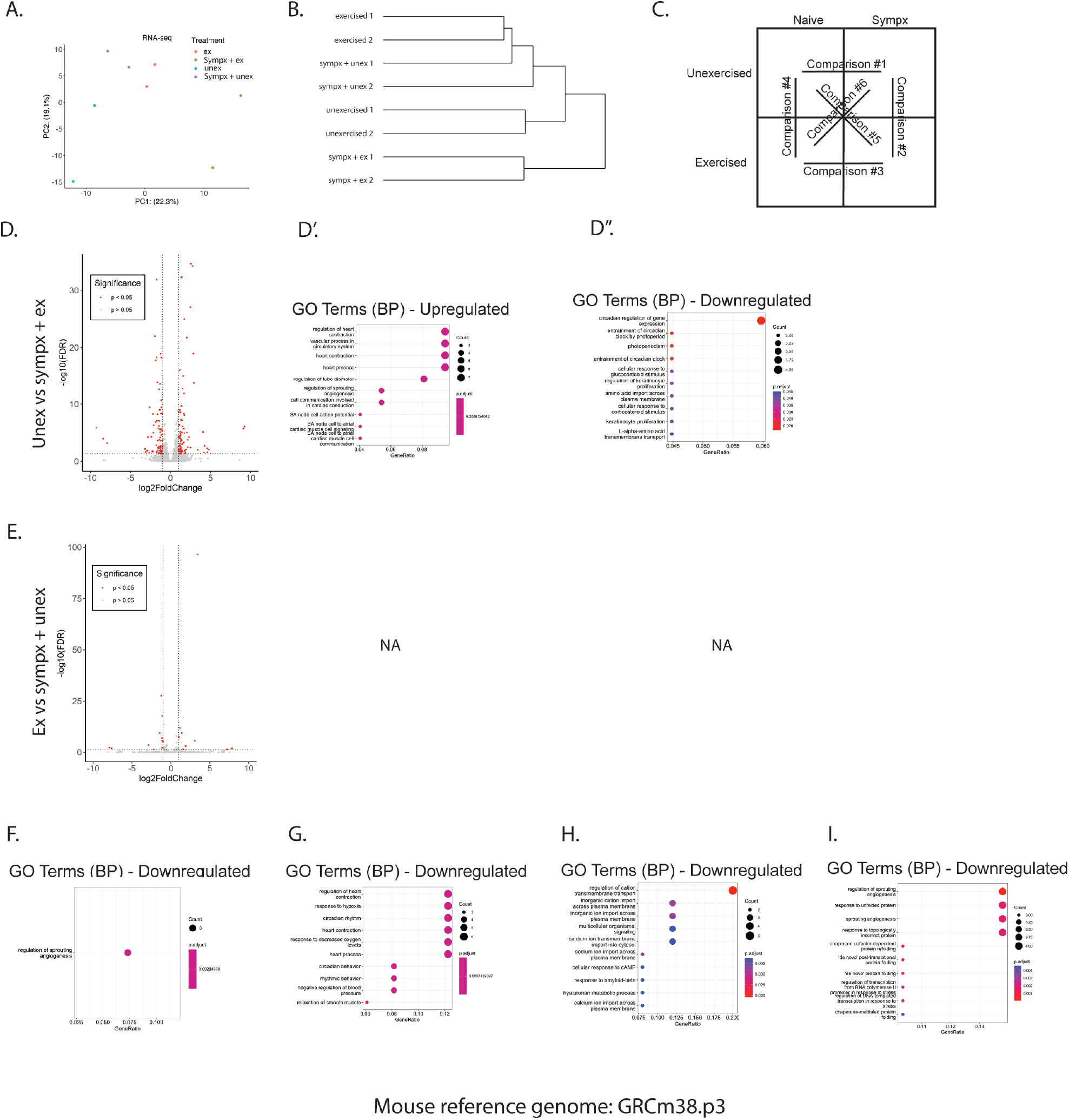
Bulk RNA sequencing of EDL muscle. **A.** PCA analysis showing clustering of different experimental groups (unex, ex, sympx + unex, sympx + ex). **B**. Hierarchical clustering of PCA analysis. **C**. Box diagram representing the 6 comparisons performed. **D-E**. RNA-sequencing results on EDL visualized as volcano plots showing up (right) and down (left) regulated genes for each comparison: D. unex vs sympx + ex E. ex vs sympx + sed. **D’-D”**. Upregulated (D’) and downregulated (D”) genes from comparison D grouped into Gene Ontology (GO) biological process (BP) pathways. F-**I**. Downregulated genes from each comparison from **Figure 4A-D** grouped into Gene Ontology (GO) biological process (BP) pathways: A. sympx + unex vs unex, B. sympx + ex vs sympx + unex, C. sympx + ex vs ex, D. Ex vs unex.

**Supplemental Figure 4.**
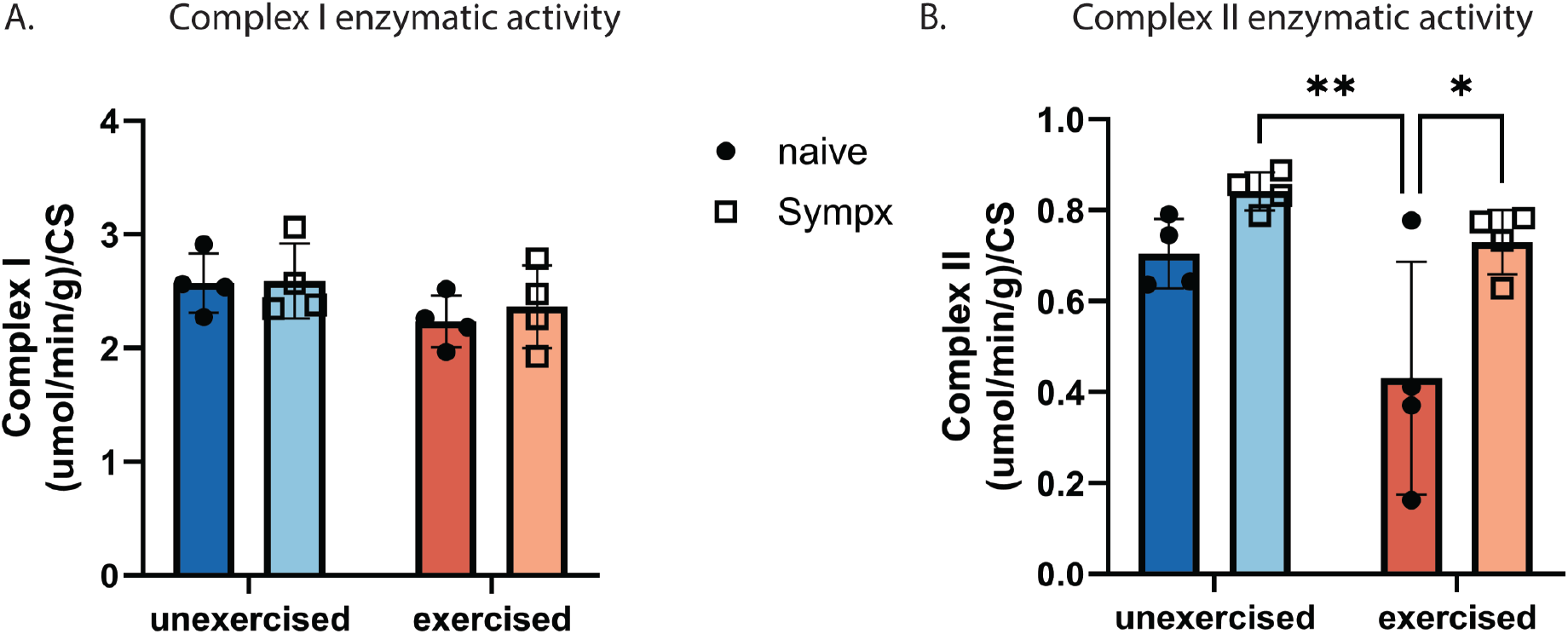
Electron transport chain enzyme assays. **A-B.** Complex 1 (A) and 2 (B) enzymatic activity in EDL of unex, ex, sympx + unex, and sympx + ex mice measured in umol/min/g and normalized to citrate synthase activity. **D**. Muscles were fed succinate dehydrogenase to determine the activity of complex II in the electron transport chain, which was similar for all four groups.

**Supplemental Figure 5.**
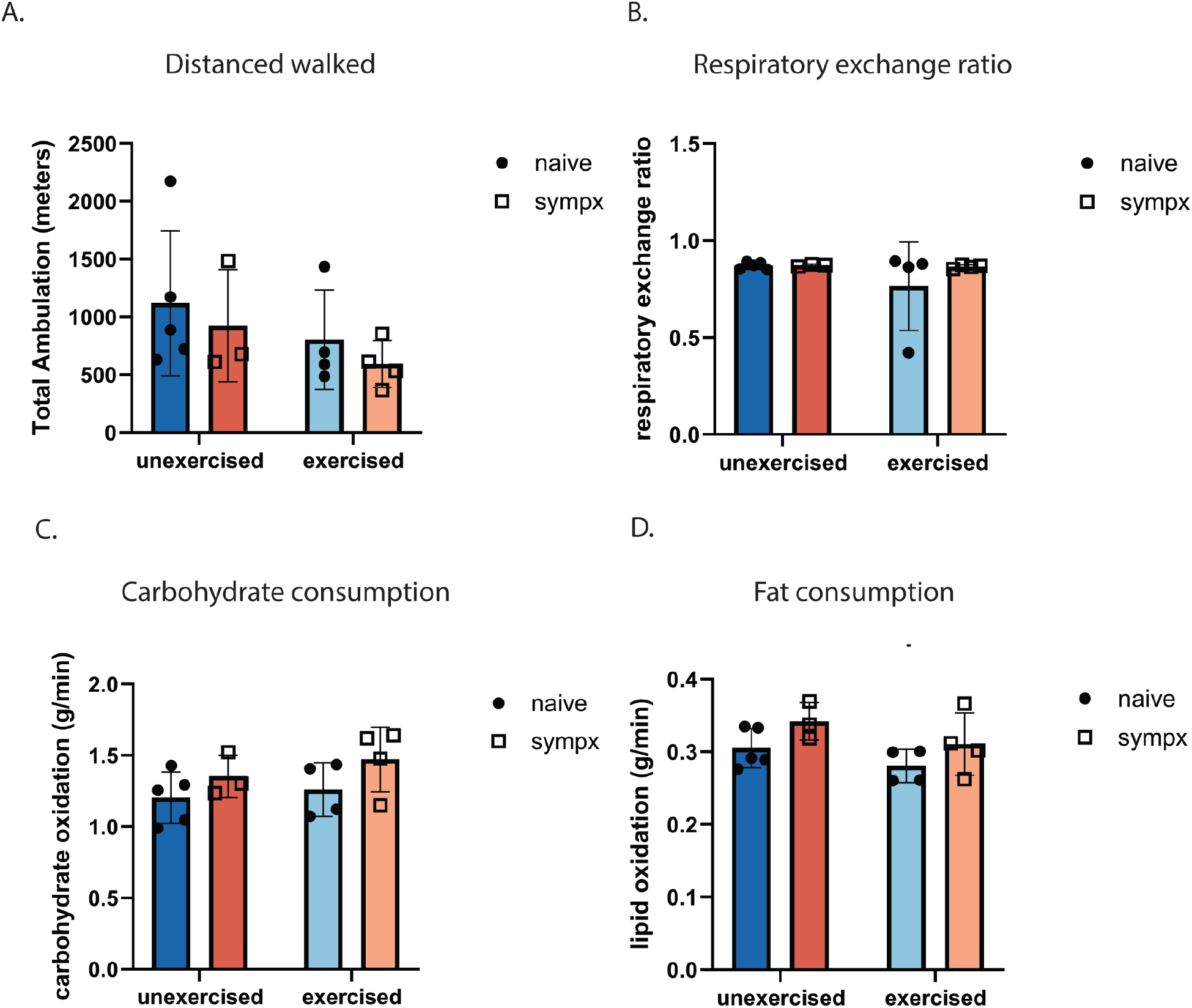
Whole-body metabolic data. Whole-body metabolic data was collected over 72 hours for unex, ex, sympx + ex, and sympx + unex mice. **A**. Total ambulation distance was measured in meters. **B**. Respiratory exchange ratio (RER) was calculated by the volume ratio of CO2 and O2. Data was collected every 5 minutes and the average RER over 72 hours was recorded **C-D**. Carbohydrate (C) and fat (D) consumption data was collected every 5 minutes and the average carb and fat consumption rate over 72 hours was recorded. A 2-way ANOVA with Tukey’s post-hoc analysis was performed for all quantifications, no significance was found for any of the comparisons.

## Acknowledgements

This work was supported by grants from the National Institutes of Health, National Institute of Neurological Disorders and Stroke award number K01NS124912 and in part by a developmental grant from the NIH-funded Emory Specialized Center of Research Excellence in Sex Differences U54AG062334 and T32GM008490. We thank Brittney Ward, Nikki Boon, and Kate Pollack for their contributions to this paper and Dr. Daniel Mucida for providing the *Adrb2* ^fl/fl^ mice.

Conceptualization, J.O. and P.J.W.; Investigation, J.O., H.S., J.H., T.K., T.T., G.W.C. Writing – Original Draft, J.O. and P.J.W.; Writing – Review and Editing, all authors; Funding Acquisition, J.O. and P.J.W.; Resources and Supervision, J.C., A.H., P.J.W.

The authors declare no competing interests.

---

**Table 1. Treadmill anaerobic exercise protocol for mice.** 1 hour exercise protocol for mice.

**Table 2. Treadmill aerobic exercise protocol for mice.** 17-day exercise protocol for mice. In addition to these durations, training will always start with a 5-minute warm up at a velocity of 10 m/min.

